# Transient effect of mossy fiber stimulation on spatial firing of CA3 neurons

**DOI:** 10.1101/324699

**Authors:** Joonyeup Lee, Miru Yun, Eunjae Cho, Jong Won Lee, Doyun Lee, Min Whan Jung

## Abstract

Strong hippocampal mossy fiber synapses are thought to function as detonators, imposing ‘teaching’ signals onto CA3 neurons during new memory formation. For an empirical test of this long-standing view, we examined effects of stimulating mossy fibers on spatial firing of CA3 neurons in freely-moving mice. We found that optogenetic stimulation of mossy fibers can alter CA3 spatial firing, but their effects are only transient. Spatially restricted mossy fiber stimulation, either congruent or incongruent with CA3 place fields, was more likely to suppress than enhance CA3 neuronal activity. Also, changes in spatial firing induced by optogenetic stimulation reverted immediately upon stimulation termination, leaving CA3 place fields unaltered. Our results do not support the traditional view that mossy fibers impose teaching signals onto CA3 network, and show robustness of established CA3 spatial representations.

## Introduction

The recurrent connectivity of the CA3 subregion of the hippocampus has led to a theoretical framework in which multiple neuronal representations are stored within the same neural network and each representation is retrieved by partial inputs [1]. A major source of external sensory information to CA3 is layer II neurons in the entorhinal cortex (EC). The same EC neurons also project to granule cells in the dentate gyrus (DG), which in turn project to CA3 via mossy fibers [2]. Mossy fibers make connections with CA3 pyramidal neurons via specialized synapses that are located at the perisomatic region, large in size with multiple neurotransmitter release sites, and also highly facilitating [3]. These characteristics would allow individual mossy fiber synapses to effectively trigger postsynaptic spikes. Due to these anatomically and functionally unique properties, mossy fiber inputs have long been considered to function as detonators (unconditional activators of postsynaptic CA3 neurons) and convey ‘teaching’ signals for setting up new neuronal representations in CA3 during memory acquisition. A new pattern of CA3 neuronal activity imposed by mossy fiber inputs is stabilized by long-term potentiation of recurrent collateral synapses in CA3 [1, 4, 5].

For an empirical test of the proposed role of mossy fibers in carrying teaching signals, we optically stimulated a subset of dentate granule cells expressing channelrhodopsin while recording CA3 neuronal activity in freely-behaving mice. We tested effects of mossy fiber stimulation on spatial firing of CA3 pyramidal neurons at two different frequencies and under two different behavioral states. We found that mossy fiber stimulation induces only transient changes in CA3 spatial firing under all conditions tested. Our results do not support the view that mossy fibers impose new patterns to learn onto CA3 network and also reveal robustness of established spatial representations in the CA3 network.

## Results

### Optogenetic stimulation of hippocampal mossy fiber

To stimulate hippocampal mossy fibers in freely-moving animals, we infected a subset of dentate granule cells with an adeno-associated virus containing a channelrhodopsin variant (ChETA-eYFP [6], Fig 1A), which is more reliably stimulated at higher frequencies than wild type channelrhopdopsin-2. eYFP expression was restricted to ~2.6% of granule cells that were distributed both in the superior and inferior blades of the dorsal dentate gyrus and mostly located near the outer border of the granule cell layer where mature granule cells are present [7] (Fig 1A and 1B). Considering that ~10% of granule cells are active in a given environment and the size of place fields is roughly 10% of the environment size, ~1% of granule cells would normally be active at a given position [8]. Thus, our stimulation is expected to activate at most ~2-3 times more mossy fibers in addition to the physiologically active ones.

**Fig 1.**
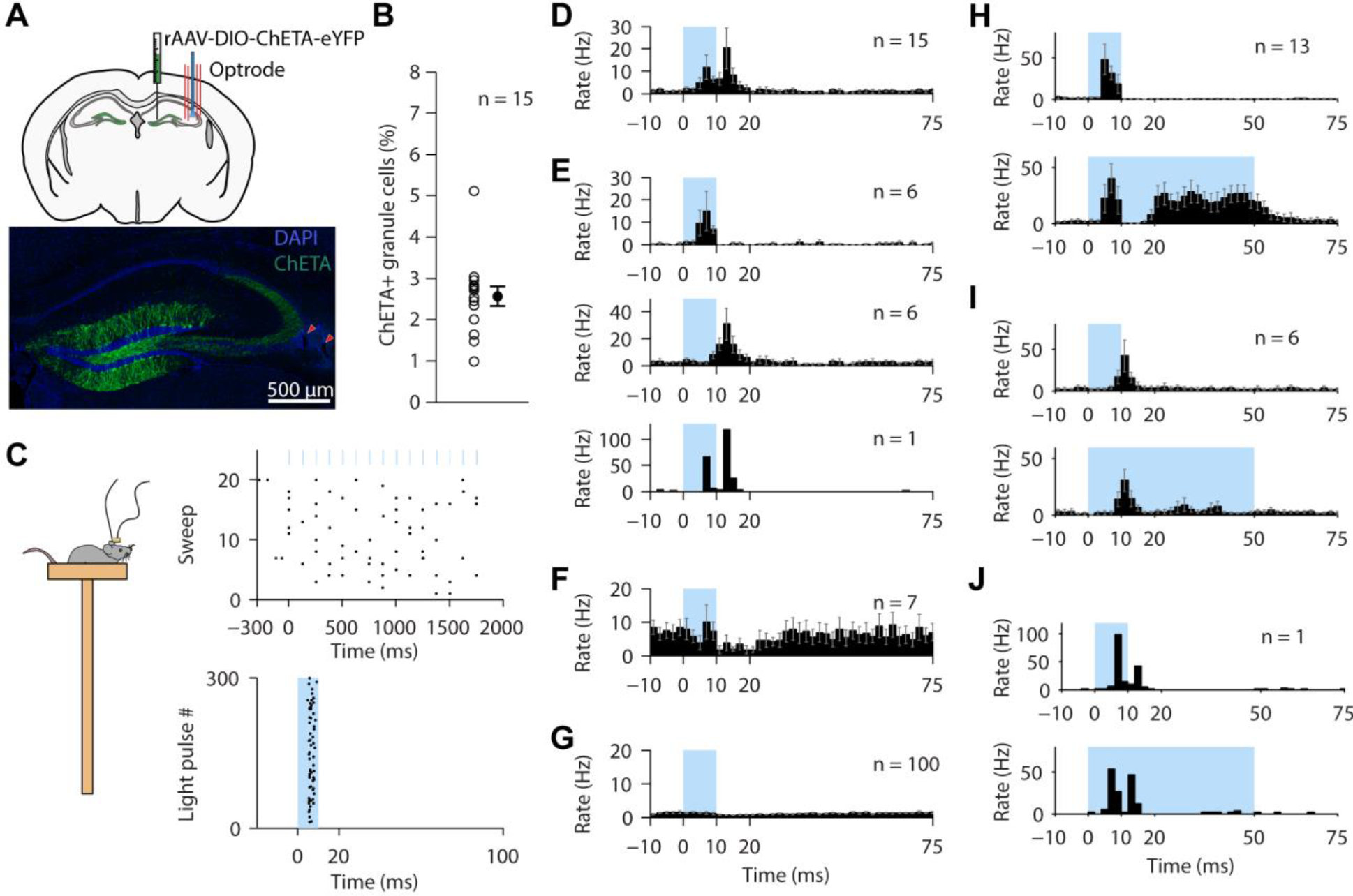
Optogenetic stimulation of hippocampal mossy fibers elicits spikes in CA3 pyramidal neurons. (A) Virus injection into the hilus and optrode implantation in CA3 (top). ChETA-eYFP (green) expression in the dentate gyrus (bottom). Red arrow heads, tracks of tetrode insertion. (B) Proportion of ChETA-eYFP expressing granule cells in each mouse (open circles) and their mean (a filled circle). (C-G) CA3 pyramidal neuronal responses to 8 Hz optogenetic stimulation of mossy fibers while mice were placed on a small platform. (C) Light-evoked spikes (black dots) of an example CA3 neuron without (top) or with (bottom) aligning to light stimulation (blue). (D) Peri-stimulus time histograms (PSTHs, 2 ms bin) for all light-activated pyramidal neurons at 8 Hz stimulation. (E) PSTHs of directly activated (top), indirectly activated (middle), and double-peaked neurons (bottom) from (D). (F) PSTH of inhibited neurons. (G) PSTH of non-responding neurons. (H-J) PSTHs of directly (H) and indirectly (I) activated neurons as well as a neuron with double peaks (J) in response to 10-ms (top) or 50-ms (bottom) light pulses (blue) repeated at 2 Hz. Error bars, SEM.

Basic characteristics of CA3 pyramidal neuronal responses to light stimulation were assessed while the mice were allowed to move freely on a small platform (5 cm × 5 cm; Fig 1C). Light stimulus (10 ms) was delivered at diverse frequencies (1 – 50 Hz). The majority (*n* = 122 neurons from 8 mice) were tested with five different frequencies (1, 2, 8, 20 and 50 Hz; 20 trains of 15 pulses at each frequency) and 8-Hz stimulation activated the largest proportion of CA3 neurons (see Fig 2C). We therefore analyzed CA3 neuronal responses to 8-Hz light stimulation. Of 122 pyramidal neurons, 15 were activated, seven were inhibited, and 100 remained unaffected (Fig 1D-G; Wilcoxon rank-sum test; see Materials and Methods). The light-activated neuronal population showed two response peaks (7.1 and 11.4 ms, Fig 1D). The population consisted of neurons with single (either early or late) or double response peaks (Fig 1E). The early peak likely reflects monosynaptic activation (direct activation) considering the EPSP-to-spike latency of the mossy fiber synapse on CA3 pyramidal neurons [9]. The late peak presumably corresponds to spikes occurring due to light-evoked recurrent CA3 inputs (indirect activation) considering the reported delay at recurrent connections in CA3 [10]. To examine whether the latter reflects rebound activity to stimulation offset, we compared CA3 neuronal responses to 10 and 50 ms-long light stimulation on the platform (*n* = 20 from nine mice). CA3 neuronal responses to 10 and 50 ms-long light pulses were similar (Fig 1H-J) indicating that the late peak is not because of rebound activity. These results indicate that our stimulation, even at a low frequency, reliably evokes spikes in a subset of CA3 pyramidal neurons in freely-moving mice.

**Fig 2.**
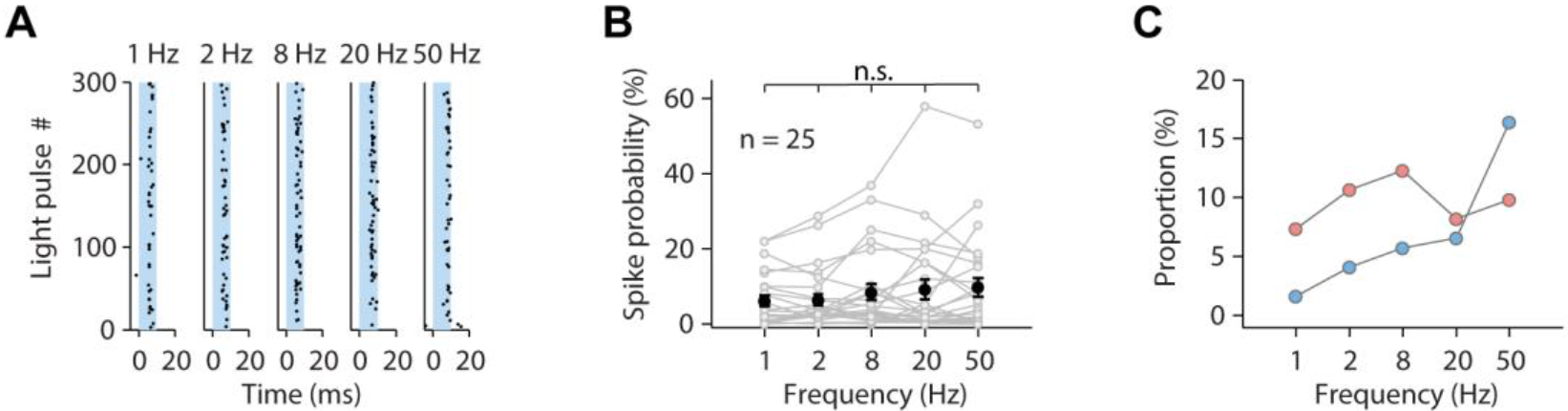
CA3 pyramidal neuronal responses to different frequencies of mossy fiber stimulation. (A) Light-evoked spikes at different stimulation frequencies for an example neuron. (B) Spike probability as a function of stimulation frequency for individual neurons that were activated by at least one frequency (open circles) and their mean (filled circles) (χ^2^(2)=2.7, p=0.615; Friedman test). (C) Proportions of light-activated (red circles) and light-inhibited (blue circles) neurons at different stimulation frequencies. Error bars, SEM.

Next, we analyzed how different stimulation frequencies affect activity of CA3 pyramidal neurons. Of the 122 CA3 pyramidal neurons tested with multiple stimulation frequencies, 25 were activated by optical stimulation at one or more stimulation frequencies. Although a few neurons showed progressive facilitation in spike probability (S1 Fig), the overall spike probability of the light-activated neurons (*n* = 25) was largely indifferent to stimulation frequency (Fig 2A and 2B). Instead, the proportion of light-inhibited neurons, but not light-activated neurons, increased with stimulation frequency, suggesting mossy fiber stimulation at higher frequencies recruits more feedforward inhibition (Fig 2C).

### Transient changes in spatial firing of CA3 pyramidal neurons upon mossy fiber stimulation

We next asked how mossy fiber stimulation affects firing of CA3 pyramidal neurons in mice navigating on a circular track for water reward at two locations for a total of 90 laps (Fig 3A). We delivered a train of either 8- or 50-Hz light pulses (duration, 10 ms) when mice (8 Hz, *n* = 15; 50 Hz, *n* = 4) were either in a running zone (running-zone stimulation sessions) or a reward zone (reward-zone stimulation sessions) during the middle 30 laps (STIM) (Fig 3A). Thus, we tested four different combinations of stimulation frequencies (8 and 50 Hz) and behavioral states (running and consuming reward). The resulting neural activity was compared to that occurring during the preceding (PRE) and subsequent (POST) 30 laps. Note that the duration of light stimulation varied across laps because we applied light stimulation while the animal’s spatial position was in the intended stimulation zone. The light stimulation did not influence the animal’s running speed in either running-zone-stimulation or reward-zone-stimulation sessions (data not shown).

**Fig 3.**
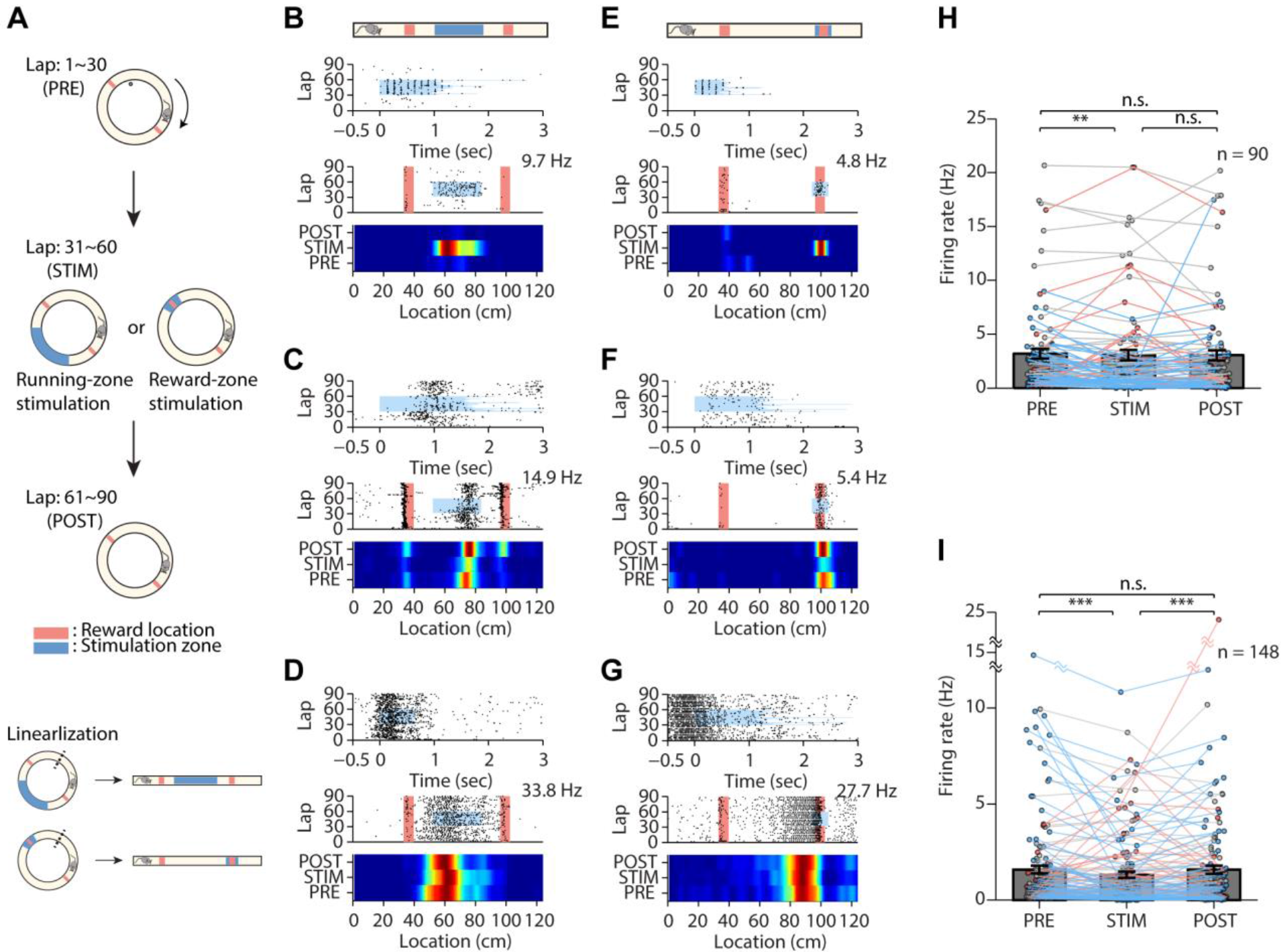
Mossy fiber stimulation at 8 Hz transiently affects CA3 neural activity during spatial navigation. (A) Schematic of experimental design. Mice navigate on a circular track for water reward at two opposite locations (red bars) for 90 laps. Light was delivered during the middle 30 (STIM) laps. The track was linearized for analysis. (B-D) Examples of light-activated (B), light-inhibited (C), and light-insensitive (D) neurons during running-zone stimulation sessions. Top, temporal spike raster plot; middle, spatial spike raster plot; bottom, heat map of spatial firing rate. Numbers on right indicate peak firing rates for the heat maps. (E-G) Examples of light-activated (E), light-inhibited (F), and light-insensitive (G) neurons during reward-zone stimulation sessions. (H,I) Firing rates for spontaneously active neurons before (PRE), during (STIM) and after (POST) light stimulation in the running zone (Friedman test: χ^2^(2)=9.2, p=0.010; post hoc Fisher’s LSD test: PRE versus STIM, **p=0.003; PRE versus POST, p=0.053; STIM versus POST, p=0.297) (H) and reward zone (χ^2^(2)=29.6, p=3.7×10^−7^; PRE versus STIM, ***p=8.9×10^−8^; PRE versus POST, p=0.072; STIM versus POST, ***p =3.9×10^−4^) (I). Inhibited (blue circles), activated (red circles) and non-responding neurons (gray circles). Error bars, SEM.

We compared mean spike discharge rates within the stimulation zone across PRE, STIM and POST blocks to examine effects of mossy fiber stimulation on CA3 neural activity during the task. With 8-Hz running zone stimulation (177 CA3 pyramidal neurons), 23 neurons (13%) significantly increased and 30 (17%) decreased their firing rates between PRE versus STIM trial blocks within the stimulation zone while the other 124 (70%) showed no significant firing rate changes (see Fig 3B-D for examples). Because decreases in firing rate can only be observed in neurons with significant baseline activity, the proportion of inhibited neurons is an underestimate. When we restricted our analysis to a spontaneously active subpopulation of neurons (see Materials and Methods), 8-Hz running zone stimulation increased firing rates in 14 neurons (16%), decreased firing rates in 30 neurons (33%) and had no impact on 46 (51%) (Fig 3H). Similar results were obtained with reward zone stimulation (182 neurons; increased: 27 (15%), decreased: 55 (30%), no change: 100 (55%); increased: 21 (14%), decreased: 55 (37%), no change: 72 (49%) with an activity threshold applied (Fig 3I; Fig 3E-G for example neurons)). The spontaneously active neuron population showed slightly, but significantly lower firing rate within the stimulation zone during STIM versus PRE blocks (Fig 3H and 3I; Friedman test followed by Fisher’s LSD test), indicating an overall inhibitory effect of 8-Hz mossy fiber stimulation on CA3 spatial firing.

Neural activity in the POST blocks recovered back to that observed during the PRE blocks (Fig 3H and 3I). In order to test how fast CA3 firing reverts back to the level before optical stimulation (PRE), we examined temporal dynamics of CA3 neural activity. Eight Hz stimulation led to various responses in CA3 firing (Fig 4A-C and 4E-G). However, altered CA3 neural activity, both increased and decreased, returned to the baseline within ~20 ms, suggesting fast attractor dynamics of CA3 network [11] (Fig 4D and 4H; cluster-based permutation test). These results indicate that the effect of 8-Hz mossy fiber stimulation on CA3 spatial firing lasts only briefly.

**Fig 4.**
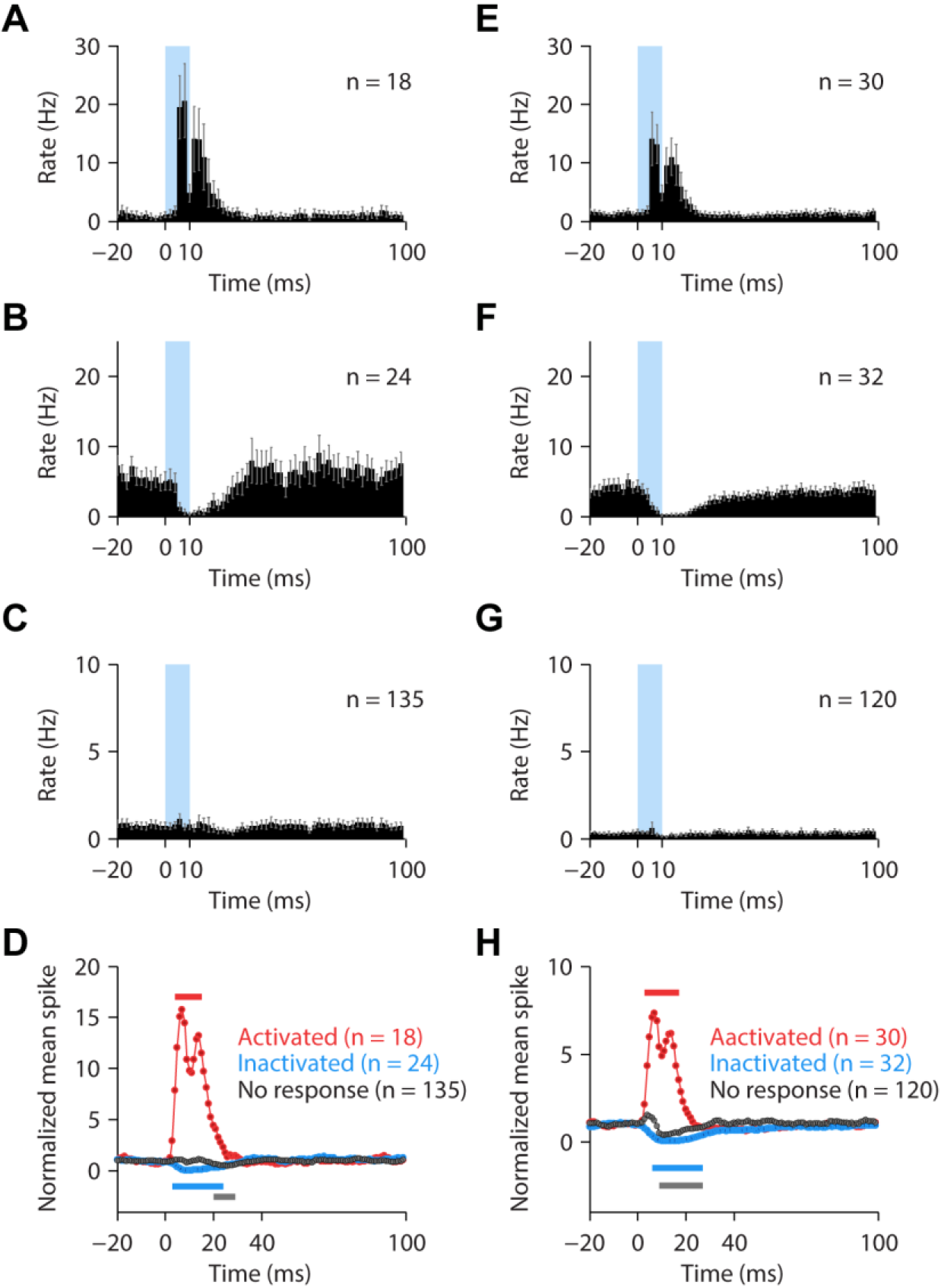
Rapid recovery of CA3 neural activity upon stimulation offset. Shown are temporal dynamics of CA3 pyramidal neuronal responses to 8 Hz stimulation during spatial navigation. (A-C) PSTHs of light-activated (A), inhibited (B), and non-responding (C) neurons recorded during running-zone stimulation sessions. Error bars, SEM. (D) Summary plots showing changes in normalized mean spike counts of activated (red), inhibited (blue) and non-responding (gray) neuronal populations. Spike counts during 1-ms time bins were smoothed with a 5-ms moving window for each neuron. The smoothed spike counts were then averaged across neurons and normalized by the mean spike count during the baseline period (20-ms time period before stimulus onset). Bars indicate the periods in which spike counts are significantly different from those during the PRE block (cluster-based permutation test, p<0.05). (E-H) Similar plots for neurons recorded in the reward-zone stimulation sessions. Note neural activity returns to the baseline (i.e., before stimulation) level within ~20 ms following stimulation offset for all types of neurons.

We obtained overall similar results with 50-Hz stimulation. Of 35 CA3 pyramidal neurons stimulated in the running zone, three neurons (8%) significantly increased and nine (26%) decreased their firing rates between PRE versus STIM trial blocks within the stimulation zone while the other 23 (66%) showed no significant firing rate changes (see Fig 5A-C for example neurons and S2 Fig for all recorded pyramidal neurons). When we restricted our analysis to a spontaneously active subpopulation of neurons, 50-Hz stimulation decreased firing rates in nine neurons (75%) and had no impact on three (25%) (Fig 5G). We made a similar observation with reward zone stimulation (46 neurons; increased: 11 (24%), decreased: 19 (42%), no change: 15 (34%); increased: four (15%), decreased: 19 (70%), no change: four (15%) with an activity threshold applied (Fig 5H; Fig 5D-F for example neurons; S3 Fig for all recorded pyramidal neurons)). The spontaneously active neuron population showed significantly lower firing rate within the stimulation zone during STIM versus PRE blocks (Fig 5G and 5H; Friedman test followed by Fisher’s LSD test), and the effect was greater compared to 8-Hz stimulation. These results show stronger inhibitory influence of 50- than 8-Hz mossy fiber stimulation on CA3 spatial firing.

**Fig 5.**
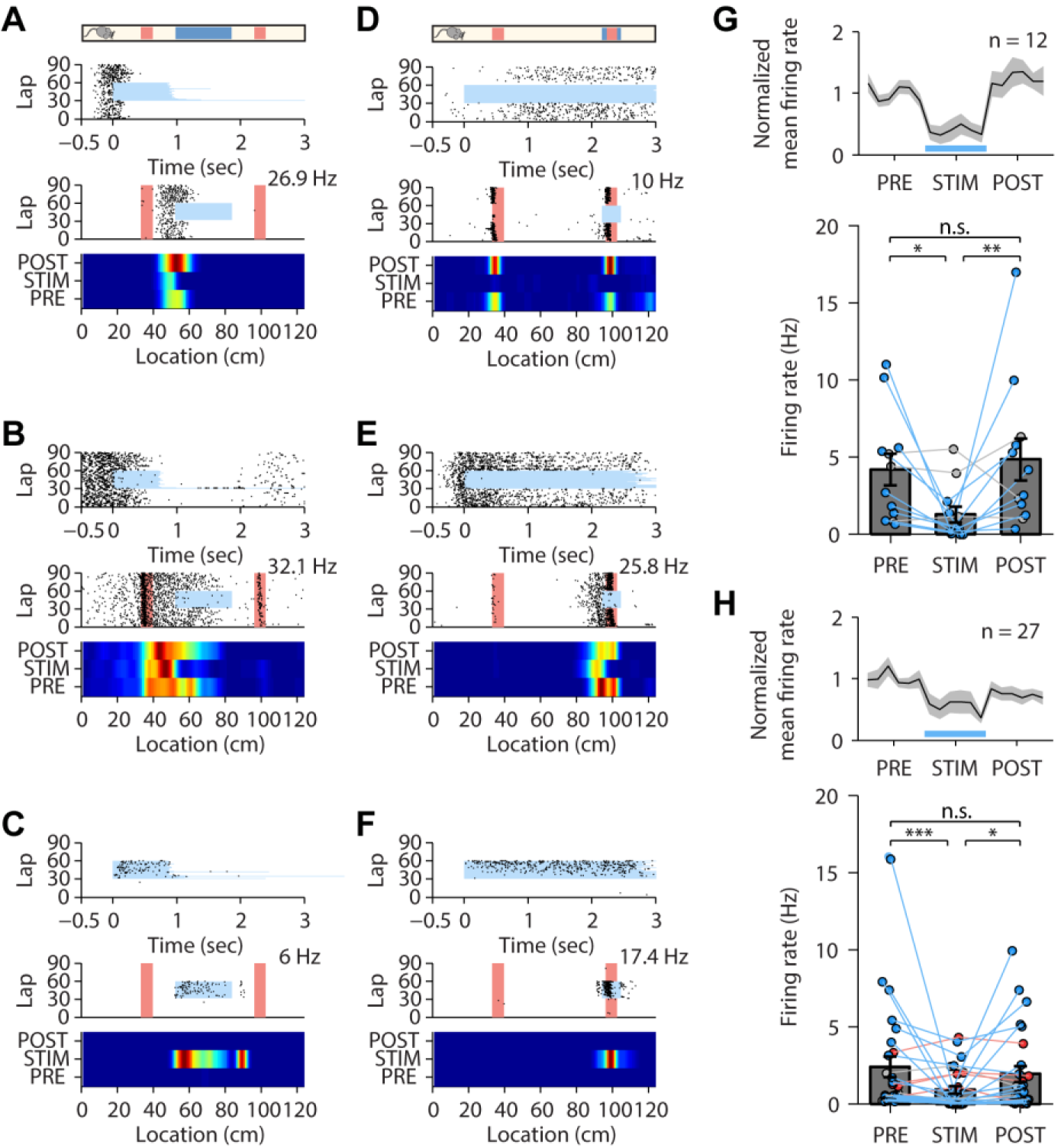
Mossy fiber stimulation at 50 Hz transiently suppresses CA3 neural activity during spatial navigation. (A-C) Examples of light-inhibited (A and B) and light-activated (C) neurons during running-zone stimulation sessions. The same format as in Figure 3. (D-F) Examples of light-inhibited (D and E) and light-activated (F) neurons during reward-zone stimulation sessions. (G and H) Normalized firing rates of spontaneously active neurons (see Materials and Methods) for running-zone (G) and reward-zone stimulation sessions (H). Top: non-overlapping five-lap mean firing rates normalized to the mean firing rate during the first 30 (PRE) laps (before the light stimulation). Bottom: firing rates before (PRE), during (STIM) and after (POST) light stimulation in running-zone (Friedman test: χ^2^(2)=8.2, p=0.017; post hoc Fisher’s LSD test: PRE versus STIM, *p=0.025; PRE versus POST, p=0.683; STIM versus POST, **p=0.008) (G) and reward-zone stimulation sessions (χ^2^(2)=13.6, p=0.001; PRE versus STIM, ***p=2.4×10^−4^; PRE versus POST, p=0.103; STIM versus POST, *p=0.041) (H). Black lines, mean; gray shading, SEM; blue bars, stimulation period. Inhibited (blue circles), activated (red circles) and non-responding neurons (gray circles). See S2 Fig and S3 Fig for all recorded neurons during 50-Hz stimulation sessions.

Although 50-Hz optogenetic stimulation altered CA3 neuronal activity within the stimulation zone for 30 consecutive laps, neural activity in the POST blocks recovered back to that observed during the PRE blocks (Fig 5G and 5H). It is difficult to perform the analysis shown in Fig 4 for the neural data obtained with 50-Hz stimulation because there are only 10-ms gaps between successive pulses (115-ms gaps in 8-Hz stimulation). Instead, we examined CA3 firing rates averaged over each consecutive five laps. The recovery of CA3 firing occurred within the first five laps in the POST blocks (Fig 5G and 5H). Collectively, these results indicate that optical stimulation of mossy fibers induces overall inhibitory and transient changes in CA3 neural activity in our task. We observed, however, exceptional cases where light-induced inhibition was sustained beyond the stimulation zone and also into the POST blocks (three neurons marked with red asterisks in S3 Fig), which may be due to facilitation of feedforward inhibitory synapses persisting beyond the stimulation period [12].

### CA3 place fields do not change after mossy fiber stimulation

Some CA3 pyramidal neurons had place fields with their centers located within the stimulation zone during the PRE block (8-Hz running zone stimulation, 34 out of 123 (28%); 8-Hz reward zone stimulation, 11 out of 89 (12%); 50-Hz running zone stimulation, 4 out of 22 (18%); 50-Hz reward zone stimulation, 5 out of 29 (17%)). Thus, the mossy fiber stimulation may be expected to strengthen the spatial information already represented by these place cells. For the majority of the CA3 pyramidal neurons, however, the same stimulation provided spatial inputs not matched by postsynaptic spatial information. Therefore, mossy fiber stimulation might be disruptive to the stability of the postsynaptic spatial map. To test how mossy fiber stimulation impacts the spatial map in CA3, we compared correlations of spatial firing across the three blocks (PRE, STIM and POST) for each CA3 pyramidal neuron (Fig 6A and 6D; see Materials and Methods). We divided the maze into two halves so that the stimulated zone is located at the center of one half (stimulated side). Spatial correlations were separately calculated for the two halves and those obtained from the unstimulated side served as a control. As an additional control, spatial correlations were also computed using those sessions in which no light stimulus was delivered (56 sessions and 130 CA3 pyramidal neurons; the half of the maze corresponding to the stimulated side was analyzed).

**Fig 6.**
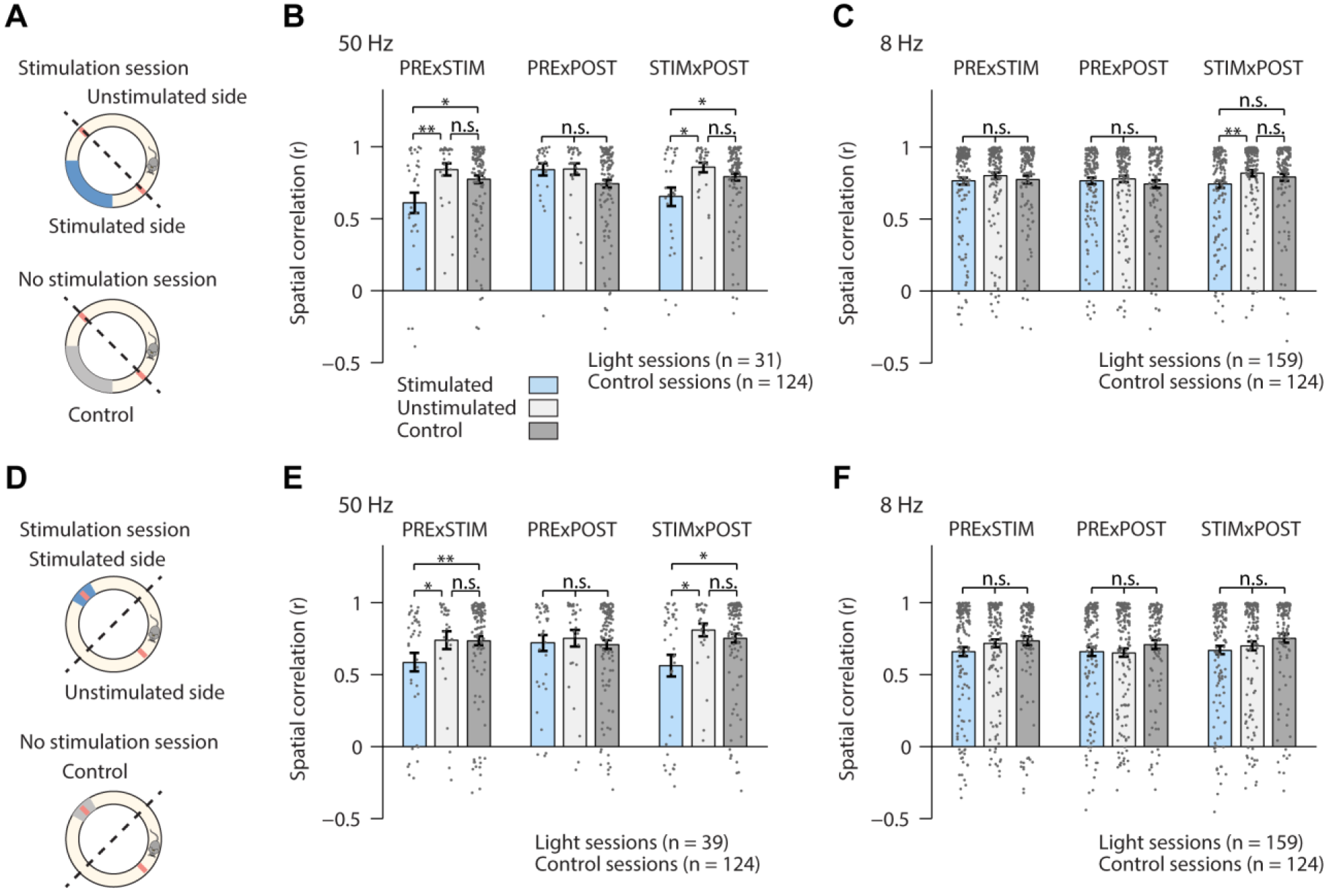
Mossy fiber stimulation at 50 Hz does not permanently disturb the CA3 spatial map. (A and D) Schematics showing how spatial correlations (r) were computed for running-zone stimulation (A) and reward-zone stimulation (D) sessions. The entire track was divided into two halves (stimulated and unstimulated). The part of the maze in no-stimulation (control) sessions corresponding to the stimulated side was also analyzed as an additional control. Pearson’s correlations between pairs of blocks (PRE, STIM, and POST) were then calculated for each neuron (B,C,E, and F). Spatial correlation values for the stimulated half, unstimulated half, and control (no stimulation) during 50 Hz running-zone stimulation (B) (Kruskal-Wallis test: PRE×STIM, χ^2^(2)=8.03, p=0.018, post-hoc Fisher’s LSD test: **p_(stimulated)×(unstimulated)_=0.005, *p_(stimulated)×(control)_=0.030; PRE×POST, χ^2^(2)=5.06, p=0.080; STIM×POST, χ^2^(2)=6.07, p=0.048, *p_(stimulated)×(unstimulated)_=0.018 and *p_(stimulated)×(control)_=0.044), 8 Hz running-zone stimulation (C) (PRE×STIM, χ^2^(2)=2.27, p=0.322; PRE×POST, χ^2^(2)=1.82, p=0.402; STIM×POST, χ^2^(2)=9.45, p=0.009, **p_(stimulated)×(unstimulated)_=0.002), 50 Hz reward-zone stimulation (E) (PRE×STIM, χ^2^(2)=8.74, p=0.013, **p_(stimulated)×(unstimulated)_=0.018 and *p_(stimulated)×(control)_=0.005; PRE×POST, χ^2^(2)=1.5, p=0.472; STIM×POST, χ^2^(2)=6.59, p=0.037, *p_(stimulated)×(unstimulated)_=0.024 and *p_(stimulated)×(control)_=0.020) and 8 Hz reward-zone stimulation sessions (F) (PRE×STIM, χ^2^(2)=2.85, p=0.24; PRE×POST, χ^2^(2)=0.515, p=0.773; STIM×POST, χ^2^(2)=4.31, p=0.116). Error bars, SEM.

With 50-Hz stimulation, PRE-STIM and STIM-POST correlations were significantly lower for the stimulated side compared to the unstimulated side and no-stimulation sessions; however, PRE-POST correlations were similar across the three groups (Fig 6B and 6E; Kruskal-Wallis test with post-hoc Fisher’s LSD test). In contrast, 8-Hz stimulation exerted no effect on spatial correlations for the stimulated side (Fig 6C and 6F). This is expected given that the effect of each light pulse on CA3 neuronal activity is only brief (~20 ms; Fig 4). CA3 spatial firing presumably depends largely on incoming sensory and internally-generated inputs except during a brief time period immediately after optogenetic stimulation. These results indicate that the effect of the mossy fiber stimulation on the spatial map is transient.

## Discussion

Strong hippocampal mossy fiber inputs, along with sparse activity of granule cells, have been proposed to drive orthogonal representations in CA3 during memory formation [1, 4]. However, it has not been tested empirically whether mossy fiber inputs are able to modify previously established neuronal representations in CA3. We stimulated mossy fibers at two different frequencies, 8 Hz and 50 Hz, while mice were navigating freely on a circular track to obtain water reward. Stimulation at 8 Hz, which is close to mean firing rates of dentate granule cells in their place field centers [8, 13], induced diverse changes in spatial firing of CA3 pyramidal neurons. However, such changes quickly (within ~20 ms) disappeared upon stimulation termination, probably due to fast attractor dynamics in CA3 recurrent network. Thus, theta-frequency stimulation of mossy fibers was ineffective in inducing long-lasting changes to established CA3 spatial representations. We also used higher frequency (50 Hz) stimulation as an attempt to maximally perturb CA3 spatial representations. Upon 50-Hz stimulation about one fourth of the place cells were partially or completely suppressed and some neurons were activated during STIM blocks. Such strong suppression or activation may be expected to produce long-term changes in CA3 place cell activity. In particular, those CA3 neurons that were activated by stimulation outside their place fields (Fig 5C and 5F) are predicted to develop new place fields that outlast stimulation, according to the conventional models [1, 4]. However, stimulation-induced firing of CA3 neurons was not maintained into the post-stimulation period. Likewise, stimulation-induced suppression of CA3 neuronal activity did not induce long-lasting changes in spatial firing of CA3 neurons, which is in contrast to CA1 place cells which become unstable and change their place field locations upon activity suppression [14]. Similar results were obtained regardless whether mossy fibers were stimulated while the animal was running toward a goal (running zone stimulation) or consuming water at a reward site (reward zone stimulation). Thus, two different patterns of mossy fiber stimulation under two different behavioral states failed to induce long-lasting changes in CA3 spatial firing in a familiar environment.

Mossy fiber activity patterns induced by our 50-Hz continuous stimulation may be considered to be relatively strong compared to naturally occurring ones considering that dentate granule cells can fire at a high frequency (> 50 Hz), but only briefly, within their place fields [13, 15] and that optogenetic stimulation presumably activates numerous mossy fibers synchronously in addition to naturally active ones. Our results show that even such strong mossy fiber inputs induce only transient changes in CA3 spatial firing. Our 8- and 50-Hz optogenetic stimulation may have been ineffective in inducing long-lasting changes in CA3 spatial firing because it was not optimally paired with innate theta oscillations in CA3. However, assuming our stimulation pulses were paired with innate theta oscillations at random phases, the stimulation would be expected to induce changes in spatial firing in a substantial portion of CA3 neurons, which was not the case. There remains a possibility that young granule cells, reported to be more excitable, exert greater impacts on CA3 pyramidal neurons [16, 17]. However, such elevated excitation is likely to be dampened by accordingly increased feedforward inhibition [18]. It is also possible that mossy fiber inputs operate in different modes according to the requirement for new memory formation. Changes in excitation-inhibition balance, by modulatory inputs for example, may allow mossy fibers to impact CA3 pyramidal neurons differently in a novel environment [19, 20], which remains to be studied.

In the conventional models, strong mossy fiber synapses, along with low divergence of dentate gyrus-CA3 projections, allow sparse granule cell activity patterns to be mapped faithfully onto CA3 pyramidal neuronal activity patterns [1, 4]. Studies in hippocampal slices and anesthetized rats have shown that single mossy fiber stimulation exerts inhibitory effects on CA3 pyramidal neurons at a low frequency, but becomes excitatory at a higher frequency (up to 50 Hz) because excitatory synaptic inputs are facilitated and exceed feedforward inhibition [21–23]. Therefore, the net effect of mossy fiber inputs largely depends on the activity patterns in the dentate gyrus, which led to a modified proposal that mossy fibers may function as ‘conditional detonators’ [21, 24]. However, optogenetic stimulation of multiple mossy fibers recruits powerful inhibitory responses lasting ~100 ms in CA3, so that activation probability of CA3 pyramidal neurons increases as a function of stimulation frequency only up to 10 Hz in anesthetized mice [25]. The effectiveness and optimal frequency of stimulation for driving CA3 pyramidal neurons may vary across studies because the ratio of excitatory to feedforward inhibitory inputs decreases as more mossy fibers are stimulated, which is supported by the finding that synchronous activity in the dentate gyrus is associated with decreased activity of CA3 pyramidal neurons [26, 27]. We also found that optogenetic stimulation of multiple mossy fibers recruits strong inhibitory responses in CA3 pyramidal neurons so that mossy fiber stimulation was more likely to suppress than enhance CA3 pyramidal neuronal activity in freely-behaving mice. Strong inhibitory responses prevented even strong (50 Hz) stimulation from unconditionally activating CA3 neurons, suggesting that granule cell activity patterns are not directly mapped onto CA3 pyramidal neuronal activity patterns in our experimental conditions. One might argue that mossy fibers might selectively activate a subset of postsynaptic CA3 pyramidal neurons in imposing a new pattern to learn. However, this argument necessarily brings excitatory/inhibitory synaptic processes in CA3 as an important player in determining which CA3 pyramidal cells are recruited. Then mossy fiber synapses are not qualitatively different from other ordinary synapses in transmitting information to postsynaptic neurons, losing their privilege of ‘imposing’ new patterns to learn, even though they may exert relatively strong influences than other excitatory synapses onto CA3 pyramidal neurons.

Previous studies in CA1 have shown that a single episode of intracellular stimulation, that is strong enough to evoke complex spikes associated with dendritic plateau potentials, can create a new place field at the stimulation location in head-fixed mice [28, 29]. A recent study also has shown that juxtacellular stimulation, that is strong enough to induce complex spikes in CA3 or CA1 neurons, can create a new place field at the stimulation location in freely-moving mice [30].

The stimulation effects on CA3 versus CA1 neurons were not compared quantitatively in this study, hence it is unclear whether and how spatial firing stability differs between CA3 and CA1. Previous studies have shown more stable spatial firing in CA3 than CA1 over time [31, 32] and in response to ventral tegmental area stimulation [33] and reward location changes [34]. Nevertheless, this study suggests that it is possible to induce long-lasting changes in CA3 spatial firing by artificially activating CA3 pyramidal neurons [30]. Our mossy fiber stimulations may have failed to create permanent place fields because they were not as effective, even at 50 Hz, for generating sufficiently strong complex spikes due to accordingly increased feedforward inhibition. Suppression of this powerful inhibitory responses, such as during exploring a novel environment [19] (but see [35]), may enable mossy fibers to strongly activate CA3 pyramidal neurons and induce long-lasting changes in CA3 spatial firing.

Dentate granule cells provide spatially well-tuned inputs to CA3 pyramidal neurons in a familiar environment [8, 13, 36, 37] where CA3 would retrieve a previously stored spatial representation. If mossy fiber inputs do not enforce a new pattern to learn onto CA3 neurons, what would be the role of mossy fiber inputs in a familiar environment? As an alternative to the pattern separation theory, the DG has been proposed to bind spatial and nonspatial information, forming distinct spatial contexts for different environments [38–40]. According to this view, mossy fiber inputs in a familiar environment play a role of informing CA3 about the current spatial context, which can be achieved by both excitatory and inhibitory control of CA3 neuronal activity. Future studies involving the silencing of mossy fibers may provide valuable information about this matter.

## Materials and Methods

All experiments were performed according to protocols approved by the Animal Care and Use Committee of the Korea Advanced Institute of Science and Technology (Daejeon, Korea).

### Animals

Seventeen male Rbp4-Cre mice (B6.FVB(Cg)-Tg (Rbp4-cre) KL100Gsat/Mmucd, stock # 037128-UCD, MMRC; 13~20 weeks old) were used. The mice were housed individually under standard conditions with a 12-h:12-h dark/light cycle. All experiments were conducted in the dark phase.

### Viral vectors

The Cre-dependent rAAV (rAAV2/Ef1a-DIO-ChETA-eYFP (ChETA-eYPF), 3.5×10^12^ particles/ml) was purchased from the University of North Carolina (UNC) Vector Core Facilities.

### Surgery and training

All surgeries were performed under isoflurane anesthesia (3% for induction; 1-2% during surgery). The animals were placed in the stereotactic frame (Kopf Instruments). A local anesthetic (lidocaine) was applied subcutaneously before opening the scalp. Two holes were made in the skull bilaterally over the target sites (1.8 mm posterior from bregma, 1.0 mm bilateral from the midline and 2.0 mm below the dura) for virus injection. ChETA-eYFP virus (500 nl) was injected bilaterally at 50 nl/min into the target sites using a Nanofil syringe (WPI), a polyethylene tubing (Becton Dickinson), and a Legato syringe pump (KD Scientific Syringe Pumps & Dispensers). The needle was kept in place for 10 min for proper diffusion of the virus before it was withdrawn. Ketoprofen (5 mg/kg) was injected subcutaneously after the surgery. The mice were allowed to recover under an infrared light lamp until they resumed normal behavior. A week after virus injection, mice were placed on water restriction. They were kept to maintain at least 80% of free-feeding body weight. After three days of handling, the mice were exposed to a circular track (inner diameter, 30 cm; outer diameter, 40 cm) that contained infrared sensors at 12 sites (1 to 12 o’clock) and four water ports spaced equally apart. The track was located at the center of a black square enclosure with four salient visual cues on the walls. During the initial training period (5 to 7 days), water reward was available at all four water ports. As the mice gradually learned to run on the track in a clockwise direction, the reward sites were reduced to two ports located at the opposite sides of the track. Well-trained mice completed 90 laps within 25 min.

A day before optrode implantation surgery, water restriction was stopped. A hyperdrive containing an optic fiber (core diameter, 200 μm) and nine tetrodes was implanted into the hippocampus of either hemisphere (1.8 mm posterior to bregma, 2.0 mm lateral to the midline and 1.5 mm below the dura). Training resumed after mice recovered for 3 to 5 days. One of the nine tetrodes remained in the cortex and was used as a reference. The optic fiber was left at the same depth throughout the whole experiments, but the tetrodes were lowered (50 to 100 μm) after each daily recording.

### Optogenetic stimulation

Light pulses (10 ms, 8 mW) were generated by a 473 nm diode laser (Omicron Phoxx) and a shutter controlled by a custom-written software (LabView 2014). Light stimulation was delivered either within a running zone or a reward zone in a given session. Since light pulses were continually delivered as long as the animals were within a stimulation zone, the number of pulses delivered varied across laps depending on the animals’ behavior.

### Electrophysiology

Unit signals were amplified 10,000 times, band-pass filtered between 600 and 6,000 Hz, and then digitized at 32 kHz. The animal’s head position was recorded by tracking two light-emitting diodes (either blue and red, or green and red) mounted on the headstage at 30 Hz. All data were acquired by the Cheetah data acquisition system (Neuralynx) and stored on a personal computer.

### Recording procedure

Two recording sessions on the track were conducted each day. They were a running-zone stimulation session and a reward-zone stimulation session in 50 Hz stimulation experiments (four mice), and a stimulation session (either a running-zone or a reward-zone stimulation session) and a control session without light stimulation in 8 Hz stimulation experiments (*n* = 15 mice). The order of these sessions was determined in a pseudo-random manner. The first and last 30 laps (PRE and POST, respectively) were without light stimulation, and light stimuli were delivered only during the middle 30 laps (STIM) in the stimulation sessions. Additional recordings were conducted on a small platform before and after each track-running session to ensure the stability of recorded unit signals. For all recorded neurons, light stimulation responses were examined at various stimulation frequencies (1 – 50 Hz) on the platform after each track running session. The majority of neurons (*n* = 122 from 8 mice) were tested with five different stimulation frequencies (1, 2, 8, 20 and 50 Hz), and, for this, a train of 15 light pulses was repeated 20 times with 15 s intervals at each stimulation frequency. Basic characteristics of CA3 neuronal responses to optogenetic mossy fiber stimulation on the platform were analyzed using 8 Hz stimulation data (Fig 1C-G). For the analysis of stimulation frequency effect, those neurons showing significant increases in firing rate at any frequency (i.e., significant light-activated neurons) were included in the analysis (Fig 2).

### Histology

When recordings were completed, an electrolytic current (20 s, 15-25 μA, cathodal) was applied through one channel of each tetrode that detected spikes to make marking lesions. Coronal brain sections (40 μm) were prepared according to a standard histological procedure. The slices were mounted with 4’,6-diamidino-2-phenylindole (DAPI)-containing Vectashield (Vector Laboratories). The brain slices were examined under a fluorescence microscope to identify the position of optic fibers and tetrodes, and to count the number of dentate granule cells expressing ChETA-eYFP.

### Proportion of granule cells expressing ChETA-eYFP

Z-stack of fluorescence images were acquired using a confocal microscope (Zeiss, LSM 780) using a 40× water-immersion objective. The percentage of eYFP-positive cells per slice was estimated in 6-12 slices per mouse (an average of 8 slices/mouse), and these values were averaged over 15 mice. The granule cell layers were outlined as a region of interest (ROI) according to DAPI signal in each slice, and the density of granule cells was calculated for each slice by counting the number of granule cells in five 20 μm × 20 μm areas. The total number of granule cells in each slice was estimated based on the total ROI area and the estimated density of granule cells. The number of eYFP-positive cells was counted manually using imageJ software (NIH).

### Unit isolation and classification

Putative single units were isolated off-line by manual cluster cutting of various spike waveform parameters using the MClust software (A. D. Redish). Those unit clusters with no inter-spike interval < 2 ms and L-ratio < 0.10 were included in the analysis. Putative pyramidal neurons and inhibitory interneurons were identified based on mean discharge rates on the track and a spike waveform parameter (peak-valley ratio), and only putative pyramidal cells were included in the analysis. The mean firing rate and peak-valley ratio of the putative pyramidal neurons were 1.54 ± 0.11 Hz and 2.29 ± 0.08, respectively (mean ± SEM). For 50 Hz stimulation experiments, total 35 and 46 putative pyramidal neurons were recorded during running zone stimulation and reward zone stimulation sessions, respectively. Because 23 of them were recorded in both sessions, we did not pool the data from the two session types together. For 8 Hz stimulation experiments, total 118 and 91 putative pyramidal neurons were recorded during running zone stimulation and reward zone stimulation sessions, respectively.

### Effect of light stimulation

To determine light-responsive CA3 pyramidal neurons on the platform, 20 sets of baseline (300 ms before the onset of a train of 15 light stimuli) and post-stimulus (20 ms after each stimulus of the train; total 300 ms) spike counts were compared using Wilcoxon rank-sum test. Those neurons with significantly (*P* < 0.05) higher (or lower) post-stimulus than baseline spike counts were considered as light-activated (or -inhibited) neurons. The rest were considered as non-responsive neurons. Because the light-activated neuronal population showed double-peak responses (*n* = 15; Fig 1D), their responses during the early (0-9 ms) and late (9-20 ms) periods were examined separately (the duration of the baseline period was adjusted accordingly). Those neurons with significantly (Wilcoxon rank-sum test, *P* < 0.05) more spikes during the early (or late) period than the baseline period were considered as directly (or indirectly) activated neurons. Two light-activated neurons (20 ms window analysis) failed to show a significantly increased activity in any of the early and late analysis windows. The effect of stimulation frequency was examined by comparing 30 sets of post-stimulus spike counts at different stimulation frequencies.

For the effect of light stimulation during track running, the firing rates within the light-stimulation zone were compared between PRE and STIM blocks (30 laps each) using Wilcoxon rank sum test. Those neurons with significantly (*P* < 0.05) higher (or lower) firing rate during STIM compared to PRE blocks were considered as light-activated (or -inhibited) neurons. The rest were considered as non-responsive neurons. Because a decrease in firing rate can only be detected in neurons with significant baseline activity, we restricted our analysis to a subpopulation of neurons with a baseline firing rate higher than a threshold (‘spontaneously active neurons’). For this, we calculated each light-inhibited neuron’s mean firing rate during the PRE block and used the lowest value as the threshold to select active neurons. We determined the threshold firing rates separately for different stimulation conditions because the duration of light stimulation differed substantially between the two conditions. The threshold firing rates of the running-zone and reward-zone stimulation sessions were 0.66 and 0.14 Hz, respectively, in 50 Hz stimulation experiments. They were 0.18 and 0.07 Hz, respectively, in 8 Hz stimulation experiments.

### Place cell identification

Two-dimensional circular firing rate maps were transformed into one-dimensional linear maps. The position and spiking data were binned into 1 cm-wide windows, generating raw maps of spike number and occupancy. A firing rate map, the number of place fields, and spatial information per spike were calculated for each unit and for each session [41]. A firing rate map was constructed by dividing the spike map by the occupancy map and applying a Gaussian kernel (SD = 2 cm) to each spatial bin. A place field was defined that met the following criteria: (1) the peak firing rate of a place field should exceed 4 Hz; (2) the total number of spikes on the track should be at least 100; (3) a place field should be a continuous region of at least 5 cm (5 bins) in which the firing rate in each bin was above 60% of the peak firing rate in the circular track; and (4) a place field should contain at least one bin above 80% of the peak firing rate in the circular track.

### Spatial correlation

A one-dimensional spatial firing rate map was constructed at 1 cm spatial resolution, smoothed with a Gaussian kernel (SD = 2 cm), and then normalized by peak firing rate for each neuron. For those sessions with light stimulation during spatial navigation, each spatial firing rate map was divided into two halves so that one half contains a light-stimulated zone at the center (stimulated half) and the other half does not contain a light-stimulated zone (unstimulated half). For those sessions without light stimulation during spatial navigation (i.e., control sessions), the part of the spatial firing rate map corresponding to the stimulated half of the light-stimulated sessions was analyzed. Three spatial firing rate maps were constructed separately for PRE, STIM and POST blocks (30 laps each) for each neuron. Then pixel-by-pixel correlations (*r*) were calculated between pairs of blocks (PRE vs. STIM, PRE vs. POST, and STIM vs. POST) for each neuron. Those neurons with session mean firing rates < 2 Hz were excluded from the analysis.

### Statistical analysis

Matlab (2014a) (MathWorks) was used for statistical analysis. All results are expressed as mean ± SEM otherwise indicated. Wilcoxon rank-sum tests were used to examine light stimulation effects on CA3 neuronal activity on the platform and to determine light-activated and -inhibited neurons on the track. Friedman tests with post-hoc Fishers’s LSD tests were used to examine the effect of light stimulation frequency on CA3 neuronal activity on the platform and to compare CA3 neuronal activity before (PRE), during (STIM), and after (POST) light stimulation on the track. Kruskal-Wallis tests with post-hoc Fisher’s LSD tests were used to compare spatial correlations in the stimulated side, unstimulated side, and control (no light) sessions. Cluster-based permutation test (p<0.05, *n* = 10,000 permutations) was used to examine temporal dynamics of firing rate changes induced by 8 Hz light stimulation (FieldTrip [42]; Fig 4D and 4H). All tests were two-tailed, and a p<0.05 was considered as a significant statistical difference unless otherwise stated.

## Acknowledgement

We thank Albert Lee for commenting on the manuscript. This work was supported by the Research Center Program of the Institute for Basic Science (IBS-R001-D1 and IBS-R002-G1).

## Author contributions

J.L. designed and performed experiments, imaged samples, and analyzed data; J.L. prepared figures with input from D.L.; M.Y. and E.C. assisted in surgery and training; M.Y. helped with image analysis and electrophysiological recording; J.W.L., D.L. and M.W.J. helped with conceptualization; J.L., D.L. and M.W.J. wrote the manuscript; D.L. and M.W.J. supervised all aspects of the work.

